# PFS alleviates bone destruction in collagen-induced arthritis mice associated with macrophage polarization

**DOI:** 10.1101/2024.06.12.598735

**Authors:** Guangchen Sun, Chunmei Bao, Xin Sun, Yingqin Liu

**Author notes:** Correspondence should be addressed to Guangchen Sun; and Yingqin Liu. These authors contributed equally to this work (Guangchen Sun, Chunmei Bao).

## Abstract

Periploca forrestii Schltr is a clinical traditional Chinese medicine for the treatment of Rheumatoid arthritis (RA). This study aimed to investigate effects of Periploca forrestii Schltr saponin (PFS) on bone destruction and macrophage polarization in arthritis model mice and to explore its possible mechanism. Arthritis was induced in two species, BALB/c and C57BL/6 mice, by immunization with type II chicken collagen. The arthritic mice were administered PFS for four weeks. The hindfeet, blood and spleen were harvested. Molecular expression was determined by ELISA, RT-PCR and immunoblotting. PFS treatment resulted in a significant reduction in paw swelling in both species of mice. PFS also reduced cartilage destruction and infiltration of osteoclasts in BALB/c mice. Furthermore, it decreased the levels of M1 macrophage cytokines while increasing the levels of M2 macrophage cytokines in the paw and plasma. Micro-CT results in C57BL/6 mice showed that PFS attenuated microstructural damage in bone tissue. PFS inhibited CD68 and affected the expression of M1 macrophage factors such as CCL-2, TLR4, and IL-1β in the mouse paw. In addition, PFS treatment increased the M2 macrophage factor CD206 and CD163. PFS inhibits the activation of ERK, JNK, and p38 and the expression of transcription factors, including STAT3, p65, and c-Fos. PFS may modulate pleiotropic macrophage polarization and thus play an ameliorative role in bone damage, therefore PFS may be an effective alternative drug for the treatment of RA.

## Introduction

Rheumatoid arthritis (RA) is a chronic and systemic autoimmune disease that impacts almost 1% of the population worldwide. It causes advanced joint damage and mainly affects the synovial tissues, cartilage and bone [1]. This immune-mediated disease involves innate and adaptive cellular compartments and their dysregulated cytokine secretion, in addition to resident cells of the skeletal and articular compartments, such as osteoclasts, chondrocytes and stromal cells [2]. Macrophages play a key role in the innate immune function that leads to inflammation. They contribute to normal tissue homeostasis by working as antigen-presenting cells to activate adaptive immunity, monitoring for tissue damage, and promoting a pro-resolution state [3, 4].

Based on microenvironment, macrophages are believed to polarize into two major phenotypes: classical activation (M1) or alternative activation (M2). M1 macrophages are derived in vitro following monocyte-derived macrophages Th-1 cytokine interferon-γ (IFN-γ) stimulation in the presence or absence of LPS [5]. M1 macrophages highly express toll-like receptor 4 (TLR4), major histocompatibility complex class II (MHC-II), cluster of differentiation 86 (CD86), inducible nitric oxide synthase (iNOS/NOS2), and interleukin (IL)-1β, and chemokine (C-C motif) ligand 2 (CCL2) and CCL5 are associated with the M1 phenotype [2]. M1 macrophages secrete pro-inflammatory factors, such as tumor necrosis factor (TNF), IL-1β and IL-6, which play a crucial role in inflammation initiation and development [6]. Conversely, M2 macrophages function in resolution in inflammation and tissue-repair pathways and express the cell surface markers CD206, CD163, and arginase (Arg) 1. M2 macrophages are induced by Th2 factors IL-4 and/or IL-13 and express high levels of IL-10, and TGF-β [7, 8]. An increase in synovial macrophages is an early marker of rheumatic disease. In patients with RA, synovial macrophages exhibit various activation states, potentially a key mediator of joint inflammation [9, 10]. In RA, macrophages are important regulators of the innate immune response and can adopt either a pro-inflammatory (M1) or anti-inflammatory (M2) phenotype. The severity of RA disease is associated with the presence of M1 and M2 phenotypes. As technology has progressed, the complexity of this heterogeneous mixture of macrophages in the synovium during disease is being revealed [9, 11, 12]. Therapeutic strategies are available to modulate the reprogramming of pro-inflammatory M1 to an anti-inflammatory M2 phenotype or block the influx of monocytes into the inflamed joint. These strategies can be used to target various inflammatory diseases, including RA [6, 7].

Periploca forrestii Schltr is a traditional medicine that has been shown to have benefits on human health, such as anti-rheumatism, improving blood circulation, irregular menstruation, stomatitis, and mastitis. A variety of chemical constituents, including triterpenes, alkaloids, and flavonoids are involved in these activities. Periploca forrestii Schltr saponin (PFS) has been shown to display immunomodulatory, and anti-inflammatory activities. In previous studies, we found that PFS treatment histologically ameliorated bone damage in two animal models of RA [13, 14]. PFS can mitigate oxazolone-induced atopic dermatitis via modulating macrophage activation in vivo and in vitro [15]. However, the role of PFS in regulating M1 and M2 macrophage polarization in CIA remains unclear. Understanding how PFS regulates these cytokines in arthritis models may be critical for the effective treatment of RA.

In this study, we further investigated the effect of PFS on bone destruction in CIA. We also investigated how PFS regulates the polarization factors of M1 and M2 that contribute to bone injury in CIA.

## Materials and methods

### Plant Materials and Chemicals

Periploca forrestii was purchased from Guangxi Yulin Pharmaceutical Co., Ltd. (Yulin, Guangxi, China), China. Saponin of Periploca forrestii (PFS) was prepared and its main ingredients were identified according to our previous reports [15]. A voucher specimen was deposited in the herbarium of the College of Pharmacy, Guilin Medical University. All other chemicals were purchased from local companies. Antibodies for CD68 (ab53444), CD206 (ab64693), ERK1+ERK2 (ab184699), phosphorylated (p) -JNK1+JNK2+JNK3 (ab124956), JNK1+JNK2+JNK3 (ab208035), and p-p38 (ab178867) were purchased from Abcam. p-ERK1/2 (4370), p38 (9212) and p-STAT3 (Tyr705-11045) were purchased from Cell Signal Technology. NOS2 (sc-7271), CCL-2 (sc-52701), STAT3 (sc-8019) and p-p65 (sc-101753) were obtained from Santa Cruz. GAPDH (60004-1-Ig) was obtained from Proteintech Group, Inc. Wuhan Sanying. Primers for Cd163, Retnla, Mrc1, NOS2, Ptgs2, IL-1β, Ifn-ɣ, TGF-β1, and Arg1 were synthesized by the Beijing Genomics Institute. Murine ELISA kits for TNF-α, CCL-2, TL-6, TGF-β1, IL-4, IL-10, and IFN-γ were purchased from Thermo Fisher.

### Animals

Female BALB/c mice and male C57BL/6 mice 6-8 weeks old weighing 20-25 g purchased from the Animal Center of Guilin Medical College were acclimated for 1 week and randomly assigned to treatment groups. Animals were housed on a 12/12-h light/dark cycle and provided food and water ad libitum. All procedures were performed under the relevant guidelines and regulations of Guilin Medical University and the ARRIVE guidelines.

### CIA Induction

The BALB/c and C57BL/6 mice were randomly divided into three groups of eight to twelve mice each: normal control, vehicle, and PFS-treated arthritic mice. CIA was induced as described by Bao et al [13]. Briefly, chicken type II collagen (CII) (4 mg/ml CII, Sigma, USA) was emulsified in complete Freund’s adjuvant consisting of 10 mL of Freund’s incomplete adjuvant plus 40 mg of Mycobacterium tuberculosis H37RA. The CII (100 µg per animal) was injected intradermally on day 1, and a booster dose of 25 µg CII was administered by subplantar injection in the hindlimb 21 days later. CIA mice were gavaged with vehicle (water) or PFS dissolved in water at 40 mg/kg daily for 28 days, starting 7 days before arthritis induction, and doses were selected by a pilot study. Ankle joint thickness was measured twice a week for each paw using a pressure-sensitive gauge (0.01 mm accuracy, Peacock Japan). On day 42, animals were anesthetized with 5% isoflurane and sacrificed by decapitation. Ankle, paw, spleen, or blood from mice were collected immediately. Spleens from C57BL/6 were collected from all animals. The tissues were washed with PBS and weighed. Spleen weights were standardized to body weight and expressed as spleen index (g/g). Blood was centrifugation at 3000×g for 10 min and the super layer of plasma was stored at −80℃ until assayed.

### Histological evaluation

Ankle joints from BALB/c mice were harvested, fixed in 10% neutral formaldehyde, decalcified by 10% EDTA for 21 days, embedded in paraffin blocks, and cut longitudinally, using a Leica microtome, into 5μm thick sections. The sections were stained with Toluidine blue and TRAP staining [16, 17]. The fraction of positively toluidine blue in the cartilage area and TRAP stained to the total bone area was measured and calculated on stained sections using ImageJ software.

### ELISA assay

Plasma was analyzed for anti-CII IgG, IgG1, IgG2a and IgM antibodies by indirect ELISA. For cytokine detection, paws, and spleen were homogenized with lysis buffer (20 mM Tris-HCl, 150 mM NaCl, and protease inhibitor cocktail (S8820, Sigma)), supernatants were collected and ELISA was performed.

### RNA extraction and RT-PCR

The paw and spleen from C57BL/6 mice were frozen in nitrogen, and ground with a pestle, total RNA was extracted using the Tengen Total RNA Extraction Kit (Centrifugal Column). RNA purity and yield were assessed by measuring the absorbance of the RNA solution at 260 nm and 280 nm. 2 μg of total RNA was reverse transcribed into cDNA using the Rapid Reverse Transcription Kit. The cDNA was then amplified using the Tengen 2×Tak PCR Master Mix RT reaction system. Primers used for RT-PCR are listed in S1. Expression was evaluated in multiple samples for each gene to reduce processing error. Gene expression was evaluated by normalizing target gene expression to that of glyceraldehyde-3-phosphate dehydrogenase (GAPDH). Band quantification and analysis were performed using Image J software (ver. 1.49).

### Micro-computed tomography

Hindlimbs from C57BL/6 mice were harvested and fixed in 70% alcohol and then sent to Guangzhou ZhongkeKaisheng Medical Technology Co. for Micro-computed tomography (μCT) scanning. The ankle joint and paw were selected and analyzed. Bone volume/total volume (BV/TV), trabecular number (Tb.N), trabecular separation (Tb.Sp), trabecular thickness (Tb.Th), joint density (Conn.D), and degree of anisotropy (DA) were calculated using ImageJ software according to the American Society for Bone and Mineral Research guidelines (Ct.Th) [18].

### Western blotting

Paws and spleens from C57BL/6 mice were homogenized in radioimmunoprecipitation assay (RIPA) lysis solution containing a protease inhibitor (Solarbio, Beijing), and protein concentrations of the resulting lysates were determined using a BCA protein assay kit (Pierce, USA). SDS-PAGE was performed to examine the primary antibody. The membranes were then incubated with second antibodies for 1 hour at room temperature. Reactive proteins were visualized using a chemiluminescence kit following the manufacturer’s instructions.

### Statistics

Statistical analysis was performed using GraphPad Prism 9.00 data analysis software. Data were expressed as mean ± standard error of the mean (SEM), and the difference between groups was determined by one-way ANOVA followed by Turkey’s Multiple Comparison. P values less than 0.05 were considered significant.

## Results

### PFS alleviates cartilage and bone destruction

We induced collagen-induced arthritis in BALB/c mice and started PFS treatment 7 days before giving them CII subplantar. As expected, the ankle joints of the mice showed rapid acute inflammation within three to seven days of the second CII exposure, peaking around day 33. The BALB/c mice consistently developed severe swelling in the ankle, wrist joint, and digits as a result of the arthritis (Figure 1(A)). On day 42, the mice treated with PFS showed significantly reduced arthritic thickness (4.37 ± 0.35) compared to the vehicle-treated arthritic group (5.27 ± 0.64) (P<0.01). Cartilage loss was assessed using toluidine blue staining, which revealed a significant decrease in proteoglycans in the articular cartilage of the ankle joints in CIA mice. Statistical analysis showed that PFS-treated mice had increased proteoglycan accumulation in the ankle articular cartilage compared to vehicle-treated mice. Additionally, TRAP staining showed that the PFS group had fewer osteoclasts surrounding the bone erosion zone in the ankle joints compared to the vehicle group (Figure 1(C)). We also examined the effect of PFS treatment on antibody changes that contribute to bone erosion. The results showed that PFS treatment significantly increased CII-specific IgG (P<0.05 in 500 and 5000 dilutions) and decreased the levels of CII-specific IgG1 and IgG2a, which are principal elements of the Th1- or Th2-type immune responses, respectively. There was also a tendency for circulating levels of CII-specific IgM to increase after PFS administration. These results suggest that PFS treatment impacted the levels of antibodies involved in bone erosion.

**Figure 1.**
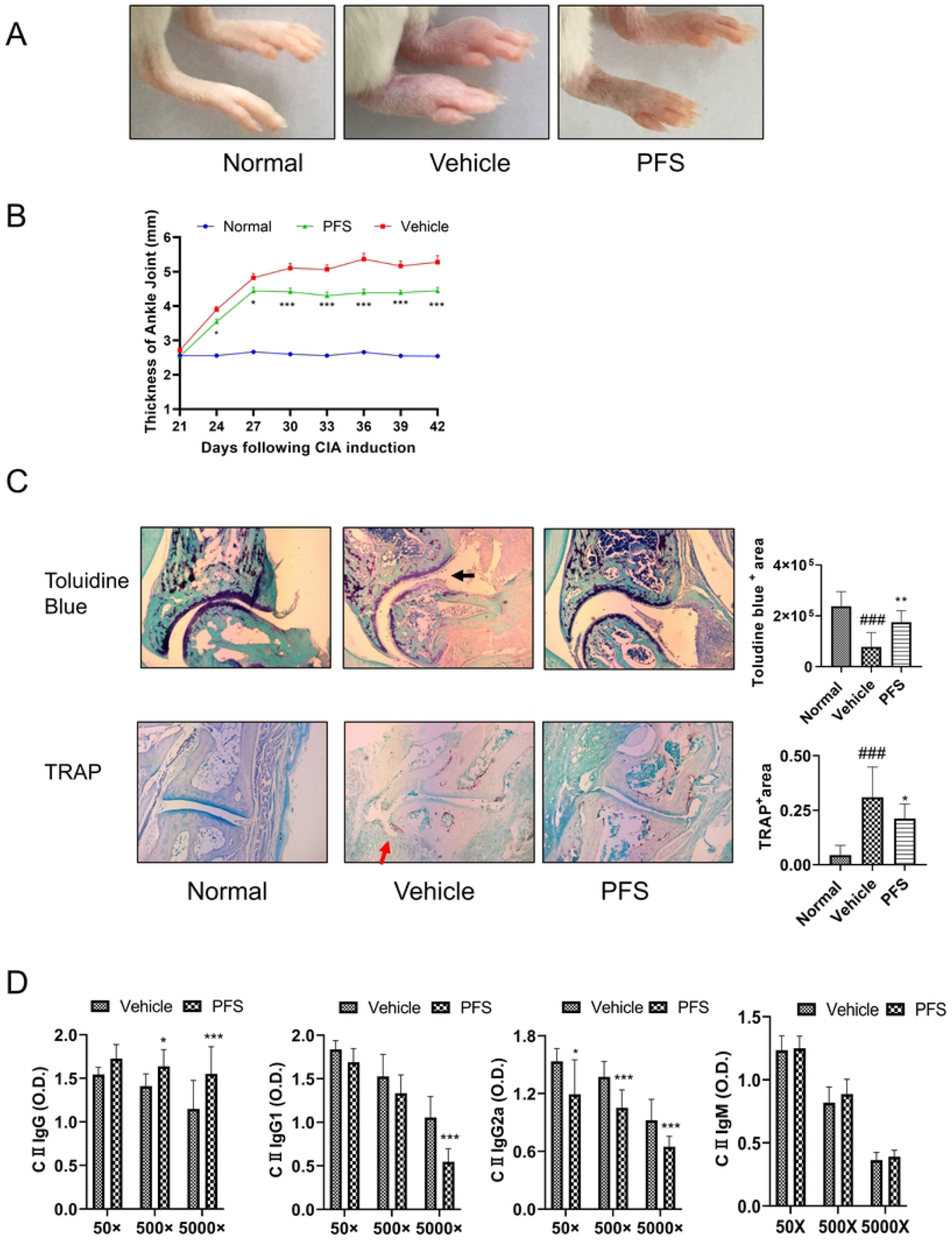
Effects of PFS on joint damage in collagen-induced arthritis in Balb/c mice. (A) Representative pictures of the hind limb on day 42. (B) Changes in mouse paw volume throughout the experiments. (C) Histological evaluation by toluidine blue and TRAP staining, black arrow: loss of proteoglycans; red arrow: osteoclast infiltrate in bone erosion site. (D) Plasma CⅡ specific-antibodies on day 42. Values represent the mean ± SD for 8-10 mice in each group. ^#^p < 0.05, ^##^p < 0.01, and ^###^p < 0.001 versus normal group, *p < 0.05, **p < 0.01, and ***p < 0.001 versus vehicle group, as determined by a one-way ANOVA followed by Tukey’s multiple comparison test.

### Effects of PFS on inflammatory factors related to macrophage polarization

The levels of inflammation factors related to macrophage polarization in the paw, plasma, and spleen of BALB/c mice were examined using an ELISA assay. According to Figure 2(A), the levels of TNF-α and CCL-2, which are M1 macrophage factors, were significantly increased in the paw tissues of vehicle-treated mice compared to normal mice. However, these effects were eliminated in PFS-treated mice. Additionally, the levels of M2 macrophage factors IL-10 and TGF-β1 were significantly decreased in BALB/c mice treated with the vehicle, but TGF-β1 levels were markedly higher upon PFS administration. PFS treatment also eliminated the elevation of plasma TNF-α levels and increased levels of IL-4, IL-10, and TGF-β1 (Figure 2(B)). Furthermore, in CIA mice treated with vehicle, there was an increase in both M1 and M2 macrophage factors in the spleen, which was reversed by PFS treatment (Figure 2(C)). These observations indicate that PFS inhibits M1 factor expression and promotes M2 factor, especially in local inflammatory sites.

**Figure 2.**
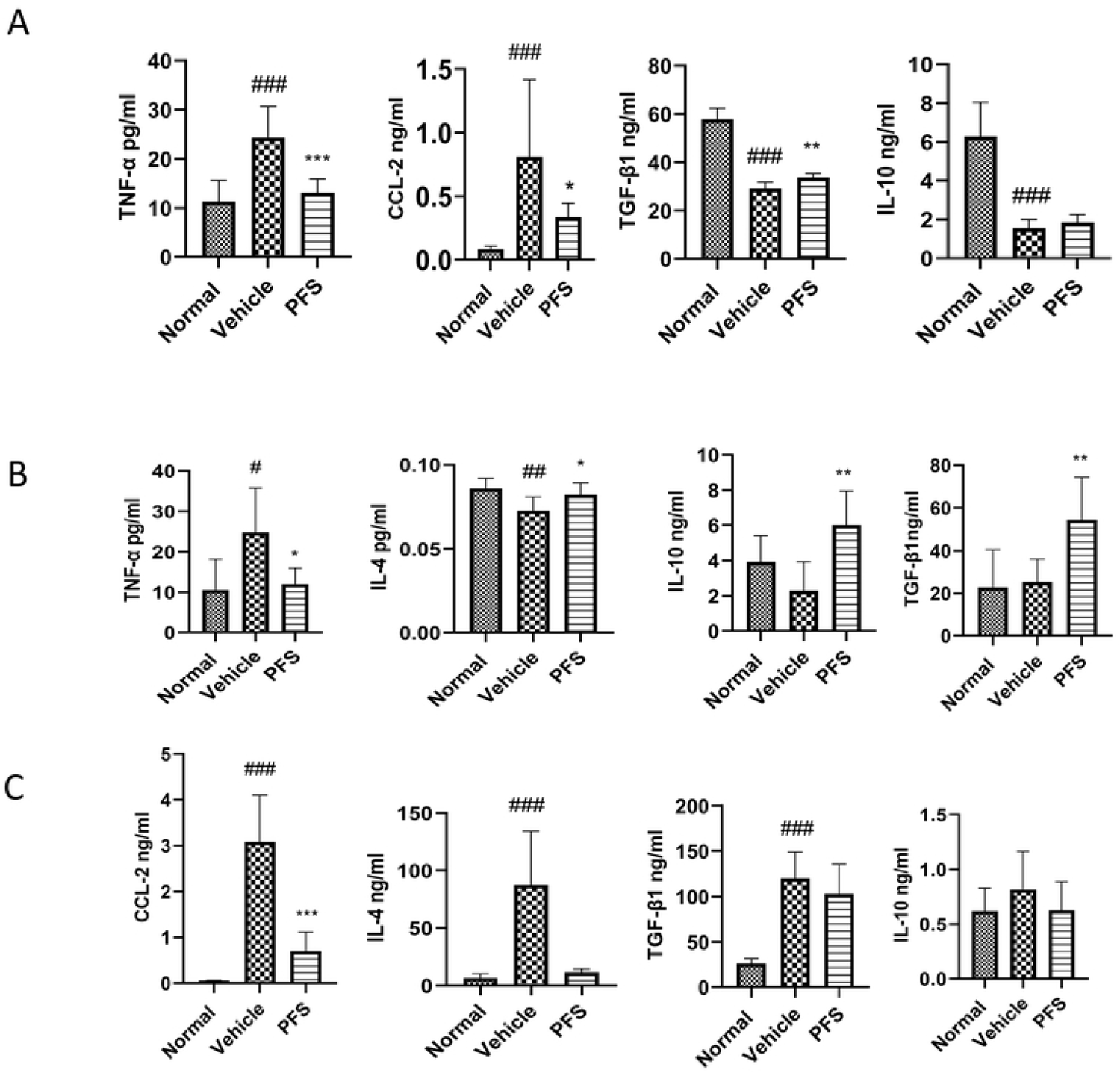
Effects of PFS on cytokine production in Balb/c mice with CIA. The cytokine levels in (A) Paw tissues, (B) Plasma and (C) Spleen. The bar graphs represent the mean ± SEM with eight to ten mice per group. #p < 0.05, ##p < 0.01, and ###p < 0.001 versus normal group, *p < 0.05, **p < 0.01, and ***p < 0.001 versus vehicle group, as determined by a one-way ANOVA followed by Tukey’s multiple comparison test.

### PFS treatment prevents bone microstructure changes

Given that PFS alleviated the inflammatory process in BALB/c mice by improving abnormal histopathological changes. The effects of PFS on CIA were examined in C57BL/6 CIA mice. PFS significantly reduced ankle swelling and spleen index with no difference in final body weight (Fig. 3(A-D)). PFS treatment significantly decreased both IgG1 and IgG2a in plasma (Figure 3(E)), and down-regulated IL-6, but up-regulated IL-10 in paws (Figure 3(F)). To evaluate bone microstructure change, we also performed μCT analyses in the hindlimb (Figure 3(G)). The mice in the normal group had intact and smooth bone surfaces in their joints. In contrast, the vehicle-treated CIA mice showed severe erosion of bone surfaces, particularly in the undivided space of the tibial joint, as well as in the tibial and fibular bones. Degenerative changes in the phalanges, such as phalangeal loss, were also evident. We then investigated whether PFS increased bone density and improved some microstructural parameters (Figure 3(H)). The BV/TV ratio was significantly lower in the vehicle-treated CIA group (0.0109 ± 0.00167%) than in the normal group (0.0189 ± 0.00284%). However, at 40 mg/kg/day in PFS treated group, BV/TV recovered significantly (0.0177±0.00294%) compared to the vehicle-treated CIA group. As expected, trabecular thickness was significantly reduced in the CIA group treated with a vehicle (0.131 ± 0.00452 mm) compared to the normal group (0.185 ± 0.00190 mm). On the other hand, PFS treatment significantly increased the thickness (0.178 ± 0.0259 mm). Simultaneously, PFS significantly increased DA and decreased connectivity. PFS treatment also resulted in higher trabecular spacing and lower trabecular number in CIA mice. These data demonstrate that treatment with PFS prevents some of the deleterious effects associated with CIA in the hindlimb bone.

**Figure 3.**
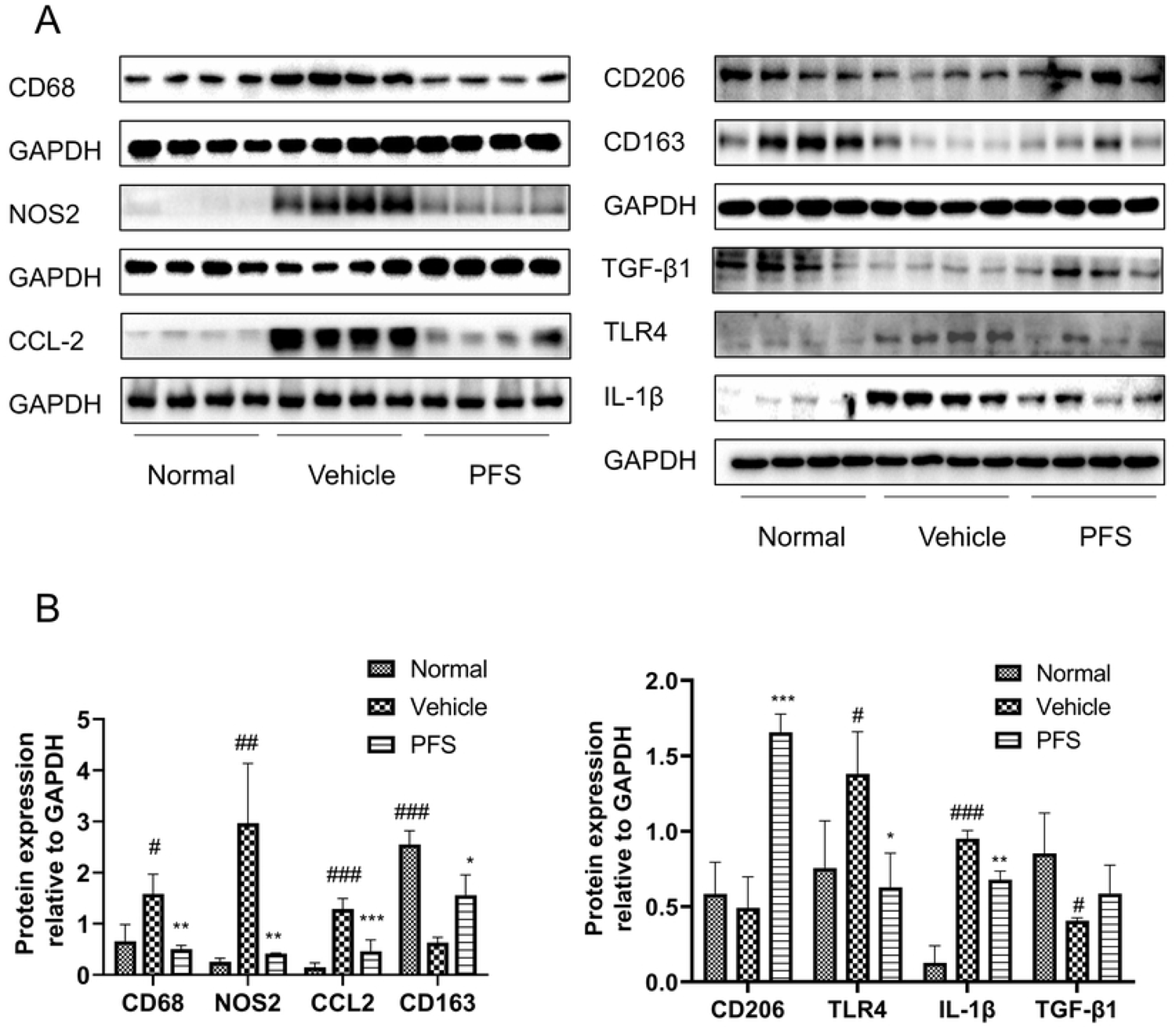
Effects of PFS on joint destruction in collagen-induced arthritis in C57BL/6 mice. (A) Representative pictures of the hind limb on day 42. (B) Changes in paw swelling throughout the experiments. (C) Body weight. (D) Spleen index. (E) Plasma antibodies. (F) Inflammatory factors. (G) Representative micro-CT pictures in the hind limb, yellow arrow: damaged and tight joint; white arrow: missing digits (H) Micro-CT parameters. The data represents the mean ± SD for 8-12 mice in each group. #p < 0.05, ##p < 0.01, and ###p < 0.001 versus normal group, *p < 0.05, **p < 0.01, and ***p < 0.001 versus vehicle group, as determined by a one-way ANOVA followed by Tukey’s multiple comparison test.

### PFS significantly regulates the macrophage polarization gene expression

The effects of PFS on gene expression associated with macrophage polarization were examined in C57BL/6 CIA mice using RT-qPCR. The bands were shown in S2. On Day 42, markedly upregulation of all M1 macrophage factors was found in the paw of the vehicle group compared to the normal group (Figure 4(A)), such as NOS2, TNF-α, IL-1β, and IL-6. Concurrently, the gene expression of all M2 phenotype markers, including CD163, Retnlα (resistin-like molecule alpha/FIZZ1), and TGFβ1 was decreased compared to the normal group. However, PFS treatment reduced a significant upregulation of all M1 marker gene expression compared to the vehicle-treated group (p < 0.05 for NOS2, TNF-α, IL-1β, and IL-6). Notably, PFS upregulated the gene expression of all M2 markers examined. In detail, PFS significantly upregulated the gene expression of CD163, Retnlα compared to the vehicle-treated group (p < 0.05 for CD163, Retnlα, and Mrc1(coding CD206); p < 0.001 for TGF-β1) (Figure 4A). Likely, mice treated with vehicle exhibited the upregulation of gene expression for primary M1 phenotype markers in the spleen, including IL-1β (p < 0.05) and Ptgs2 (coding COX2) (Figure 4(B)). Additionally, a distinct downregulation of the investigated M2 markers was observed, including Mrc1 (p < 0.01), Retnlα (p < 0.01), and TGF-β1 (p = 0.3631). Compared to the vehicle group, PFS treatment significantly upregulated the expression of M2 markers Mrc1, Retnlα, and TGF-β1, and M1 markers IL-1β and Ptgs2. Furthermore, PFS treatment was observed to reverse the expression of IFN-γ, which is associated with Th1 and activated macrophage differentiation in the paw and spleen. These results suggest that PFS has a positive effect on the regulation of macrophage polarization in the paw and restores the polarization balance of macrophage factors in the spleen.

**Figure 4.**
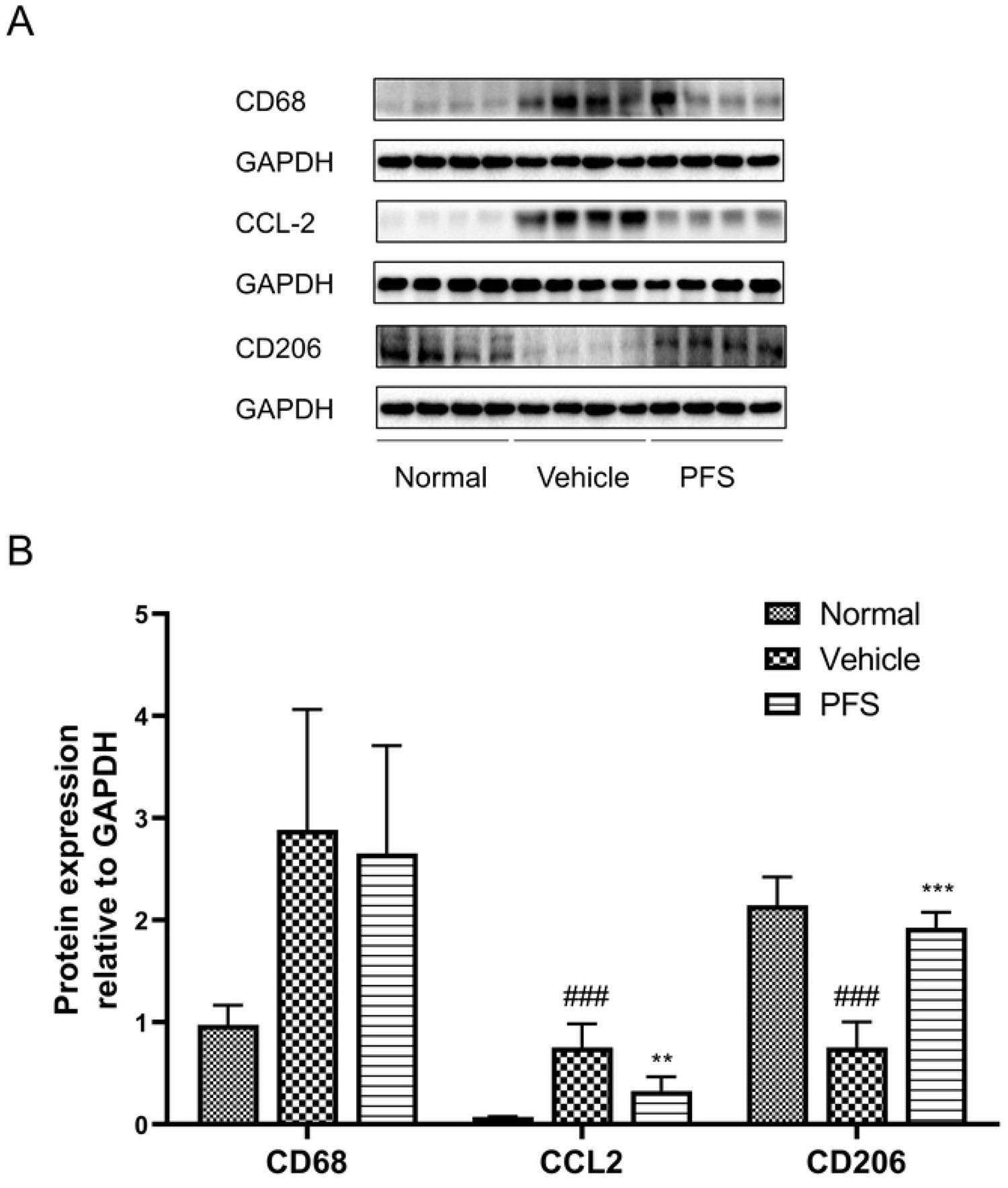
Effects of PFS on the mRNA expression of proinflammatory and anti-inflammatory factors involved in M1 and M2 polarization in C57BL/6 mice with CIA. The quantitative data of mRNA in paw tissues (A) and (B) Spleen. The bar graphs show the mean ± SEM with six to eight mice per group. #p < 0.05, ##p < 0.01, and ###p < 0.001 versus normal group, *p < 0.05, **p < 0.01, and ***p < 0.001 versus vehicle group as determined by a one-way ANOVA with Tukey’s multiple comparison test.

### PFS regulates macrophage polarization markers

The modulation of these macrophage phenotype markers in ankle joints was also investigated at the protein level in CⅡ-induced arthritis. The mice treated with vehicle showed increased expression of the M1 macrophage-related factors CD68, NOS2, and CCL-2, along with decreased expression of M2 macrophage markers CD206 and CD163 compared with the normal group. However, PFS treatment decreased the expression of M1 markers CD68 and NOS2 but increased the expression of M2 markers. Notably, PFS treatment led to a reduction in the expression of TLR4 and IL-1β, classic M1 macrophage factors, and an increase in the expression of TGF-β1, a classical M2 macrophage cytokine, in the paw of CIA mice (Figure 5). For the systemic state of macrophage polarization, we found that there was an apparent increase in M1 macrophage factor expression of CD68 and CCL-2, but a decreased expression of M2 macrophage factor CD206 in the spleen of the vehicle-treated group, but all these factors were reversed by PFS treatment (Figure 6). This confirmed that PFS inhibited macrophage polarization throughout the body, not just in the joint.

**Figure 5.**
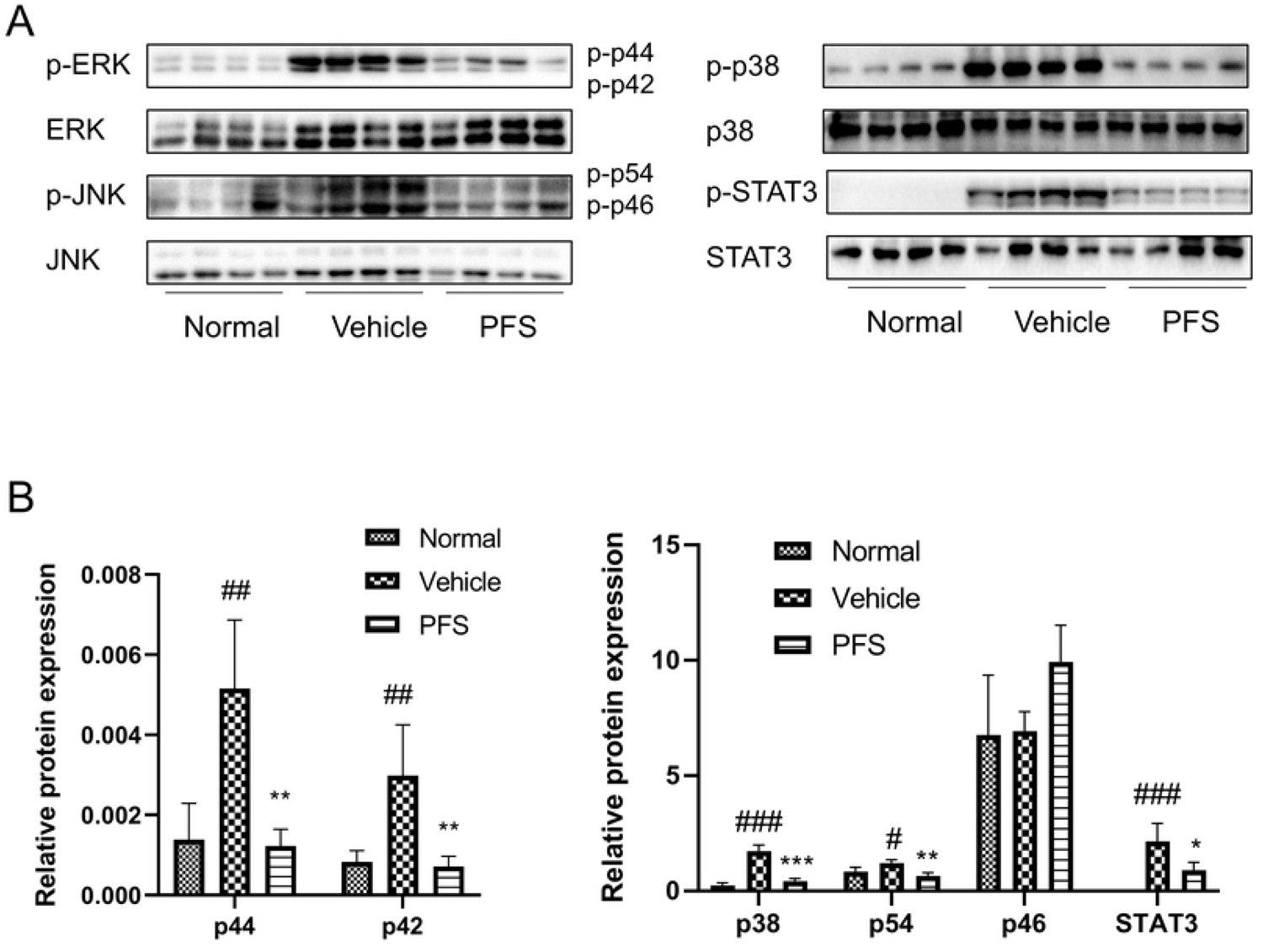
Effects of PFS on macrophage polarization factors in C57BL/6 mice with CIA. (A) Western blotting. (B) the quantitative data on target proteins. Values represent mean ± SEM with four mice per group. #p < 0.05, ##p < 0.01, and ###p < 0.001 versus normal group, *p < 0.05, **p < 0.01, and ***p < 0.001 versus vehicle group as determined by a one-way ANOVA with Tukey’s multiple comparison test.

**Figure 6.**
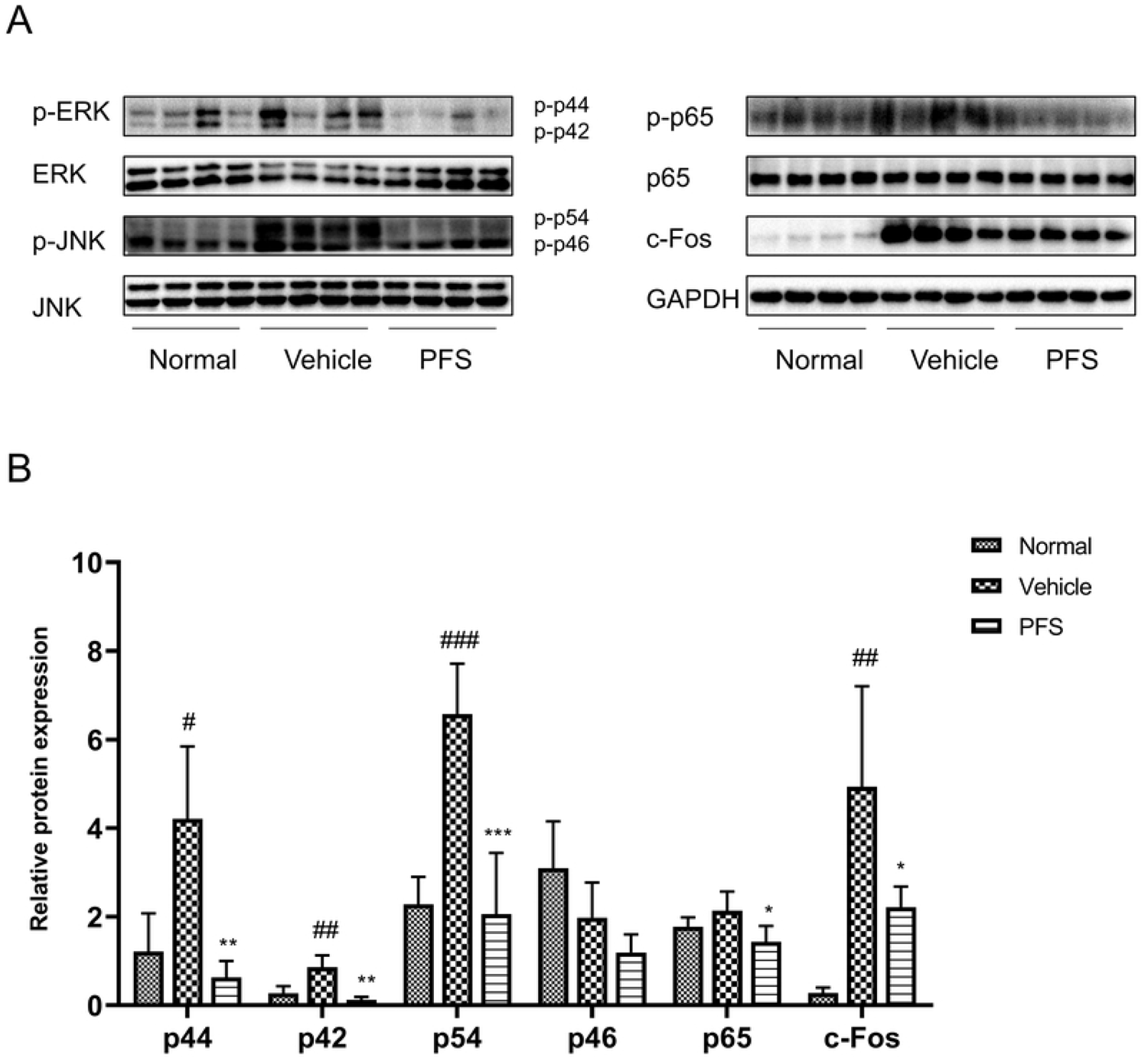
Effects of PFS on macrophage polarization factors in C57BL/6 mice with CIA. (A) Western blotting. (B) The quantitative data on target proteins. Values represent mean ± SEM with four mice per group. #p < 0.05, ##p < 0.01, and ###p < 0.001 versus normal group, *p < 0.05, **p < 0.01, and ***p < 0.001 versus vehicle group as determined by a one-way ANOVA with Tukey’s multiple comparison test.

### Effect of PFS on MAPK signaling pathway

The protein expression of the MAPK signaling pathway was analyzed using western blot analysis. The mice treated with vehicle exhibited higher levels of phosphorylated p-p38 and STAT3, as well as the phosphorylation of p44 and p42 (ERK subtypes) and p54 (JNK subtype) in the paw compared to the normal control group. However, treatment with PFS reduced the phosphorylation of p-p38, p44, p42, and p54, as well as the phosphorylation of STAT3 (Figure 7). Likewise, the systemic activation of the MAPK signaling pathway was decreased in CIA mice treated with PFS, as indicated by the reduced phosphorylation expression of p44, p42, and p54 in the spleen (Figure 8). Additionally, PFS treatment resulted in a reduction in the phosphorylation of STAT3, as well as a decrease in the expression of p65 and c-Fos in the spleen of CIA mice.

**Figure 7.**
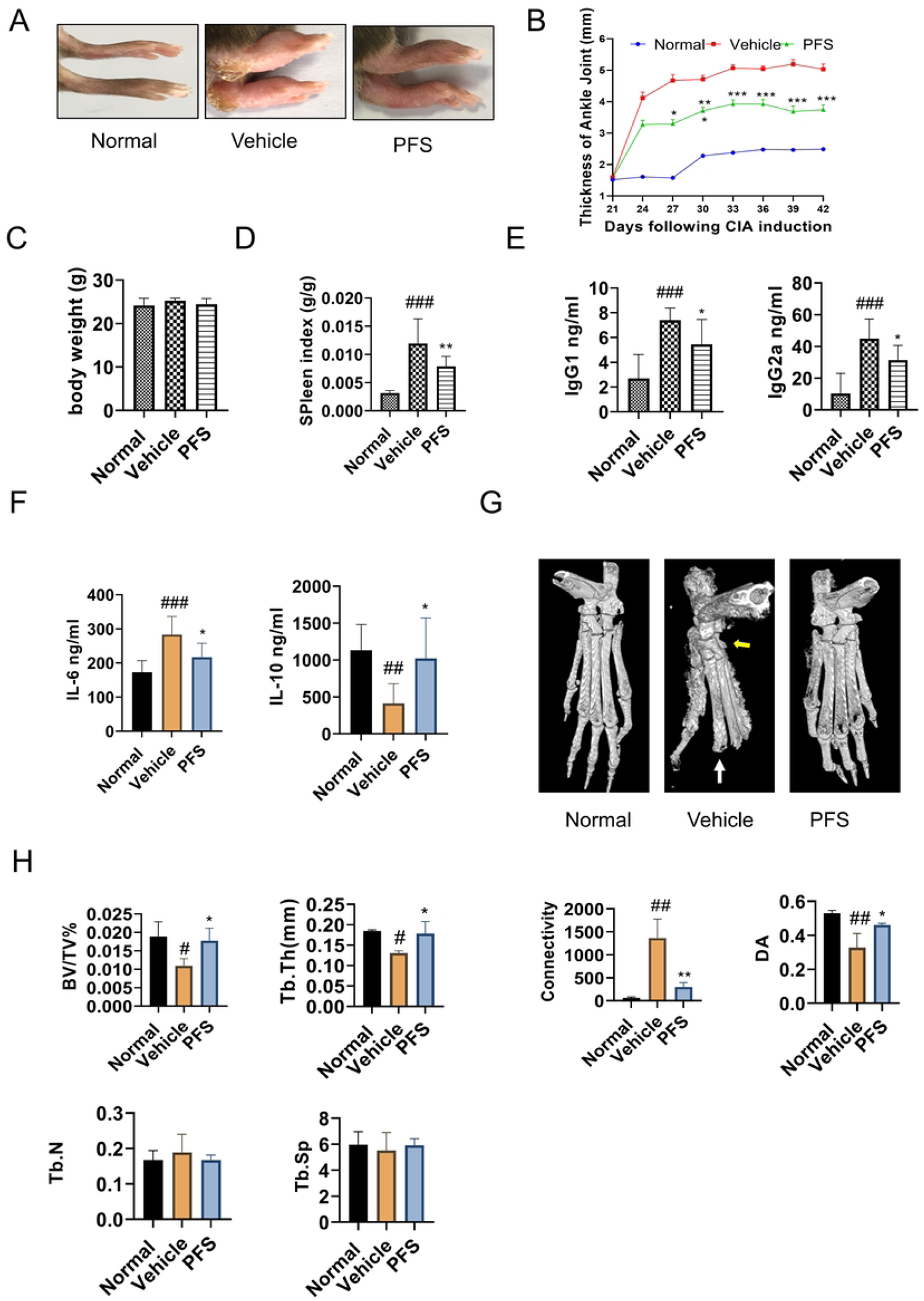
Effects of PFS on MAPK-related factors in C57BL/6 mice with CIA. (A) Western blotting. (B) The quantitative data on target proteins. Bar graphs represent the mean ± SEM with four mice per group. #p < 0.05, ##p < 0.01, and ###p < 0.001 versus normal group, *p < 0.05, **p < 0.01, and ***p < 0.001 versus vehicle group as determined by a one-way ANOVA with Tukey’s multiple comparison test.

**Figure 8.**
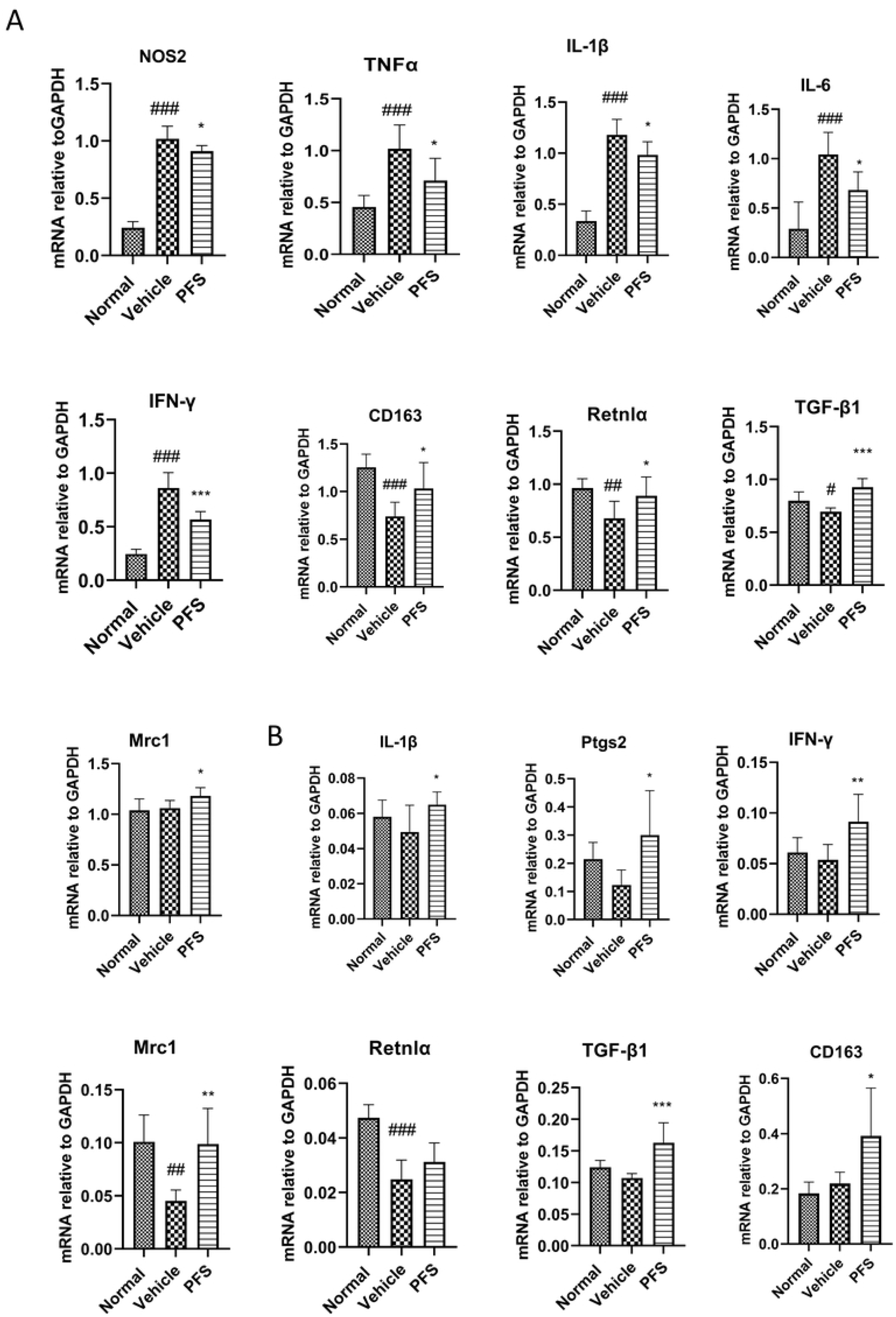
Effects of PFS on MAPK-related factors in C57BL/6 mice with CIA. (A) Western blotting. (B) The quantitative data on target proteins. Bar graphs represent the mean ± SEM with four mice per group. #p < 0.05, ##p < 0.01, and ###p < 0.001 versus normal group, *p < 0.05, **p < 0.01, and ***p < 0.001 versus vehicle group as determined by a one-way ANOVA with Tukey’s multiple comparison test.

## Discussion

Natural plants have been invaluable resources for developing new drugs, and about 80 % of drugs were either natural products or analogs by 1990 [19]. We previously found that PFS promotes the arthritis index or histological score in arthritis models, and PFS reduces the mRNA levels of inflammatory mediators of synoviocytes and splenocytes in vitro [13, 14]. The current study found that PFS attenuated articular cartilage damage and reduced osteoclast infiltration. Studies have reported that IgG and its isoforms are important in osteoclastogenesis, and anti-CII antibodies are known to be arthritogenic [20, 21]. We investigated the effect of PFS on plasma anti-CII antibody levels and found that CII-specific IgG responses were increased in CIA mice after long-term PFS treatment. Interestingly, IgG1 and IgG2a anti-CII autoantibody levels, two IgG subtypes, were significantly decreased in both CIA mice after administration of PFS. The results suggest that PFS influences CIA articular bone damage by modulating immunoglobulin IgG antibody production.

The production of inflammation-related factors was examined in CIA BALB/c mice using ELISA assay, locally (in joints and paws) and systemically (in blood and spleen). We hypothesized that the effects of PFS on CIA mice are associated with different cytokine profiles in the systemic circulation and at local sites of inflammation. The present study’s results indicate that PFS significantly reduced M1 cytokines (such as TNF-α) and increased anti-inflammatory M2 cytokines (such as IL-10 and TGF-β1) in both the paw and serum. This suggests that PFS treatment shifts the cytokine profile from M1 to M2 in the locally inflamed paw and systemic circulation of arthritic mice, thereby exerting an anti-inflammatory effect. CII induction resulted in elevated expression of both pro- and anti-inflammatory factors in the spleen, which is distinct from classical M1 and M2 cytokine polarization. However, PFS inverted the expression levels of these factors.

Many studies have revealed that macrophages are crucial for the control of bone homeostasis [22, 23]. Studies found that in active RA, high concentrations of M1-like polarized macrophages localize in the synovium [24]. They produce large amounts of pro-inflammatory cytokines iNOS, TNF-α, CCL-2, and other pro-inflammatory mediators that amplify inflammation, resulting in further damage during RA and CIA pathogenesis. TNF and IL-1 are important drivers of bone and cartilage damage in RA [25, 26]. M2 macrophages promote angiogenesis, tissue remodeling, and repair by producing high levels of anti-inflammatory factors including IL-10, TGF-β, and Arg1[6, 27]. Macrophages can also directly differentiate into mature osteoclasts influenced by the inflammatory environment, further exacerbating bone erosion [28, 29]. This is supported by osteoclastogenic macrophage subsets found in synovial tissues of mice with collagen-induced arthritis [30]. The present study demonstrated the modulatory effect of PFS on the imbalance between M1 and M2 macrophages by inhibiting the M1 macrophage chemokine CCL-2 and upregulating the M2 macrophage factors CD163 and CD206 in the paw. PFS also further downregulated the M1 macrophage pro-inflammatory mediator TLR4 and IL-1 while upregulating the M2 macrophage anti-inflammatory TGF-β1. As a result, PFS helps eliminate persistent inflammation and contributes to increased bone mass and structural integrity. Strategies that increase M2-type macrophage factors while inhibiting M1-type macrophage factors, such as inhibition of the chemokine CCL-2-induced proinflammatory M1 macrophage infiltration, are a promising approach for the safe treatment of acute and chronic inflammatory diseases [11, 31–33]. Our study demonstrated that PFS effectively modulated macrophage polarization in C57BL/6 mice, and combination with the findings of the modulatory effects of macrophage factors in BALB/c, we can infer that PFS shifted pro-inflammatory M1 macrophages to M2 macrophages, thereby inhibiting pro-inflammatory infiltration in the joints, which may be the key to the direct attenuation of CIA and amelioration of bone tissue damage.

MAPKs are a group of proteins consisting of c-Jun NH2-terminal kinase (JNK), p38 MAPK, and extracellular signal-regulated kinase (ERK). MAPKs participate in a variety of cellular activities, including proliferation, differentiation, apoptosis, survival, inflammation, and innate immunity [34, 35]. MAPKs are critical regulators of macrophage activation and proliferation [36], cytokine production [37], and monocyte migration and differentiation [33, 38]. ERK activation is associated with high expression of pro-inflammatory cytokines in macrophages [39]. JNK is required for M1 macrophage development, which promotes inflammation while inhibiting JNK polarizes macrophages to an anti-inflammatory M2 phenotype [40]. Additionally, p38 MAPK is a key signaling molecule that regulates the genesis and functions of chondrocytes, osteoblasts, and osteoclasts, which are responsible for physiological bone development and homeostasis [41, 42]. Inhibiting MAPKs has been shown to alleviate inflammation and tissue destruction in rheumatoid arthritis [43–47]. Our in vivo results confirm that PFS antagonizes the phosphorylation of the three MAPK pathway kinases. This is supported by the inhibitory effect of PFS on the transcription factors NF-κB element p65 and activator protein-1 factor c-Fos, which are activated by the MAPK pathway. This finding may be further validated in the mechanism of action of PFS on macrophage polarization and function in vitro experiments [48, 49].

## Conclusions

PFS reduced paw swelling and bone destruction, inhibited osteoclast infiltration, and regulated macrophage levels. It also attenuated microstructural damage in bone tissue and modulated macrophage polarization by inhibiting M1 macrophage factors and upregulating or reversing dysregulated M2 macrophage factors. Additionally, PFS suppressed MAPK pathway proteins and adjusted their associated transcription factors. The ability of PFS to regulate bone erosion by interfering with macrophage polarization may be a valuable tool for the treatment of inflammatory arthritis.

## Data Availability Statement

The original contributions presented in the study are included in the article/supplementary material, further inquiries can be directed to the corresponding authors.

## Ethics Statement

The animal study was reviewed and approved by Animal Experimental Ethics Committee of Guilin Medical University.

## Author Contributions

GS and YL initiated the project. GS and XS were responsible for designing experimental ideas and data analysis, protocol designing, and draft editing. CB conducted the experimental work and processed the data. GS and YL supervised and conducted the experimental work. All authors contributed to the article and approved the submitted version.

